# Dynamic chromatin remodeling in cycling human endometrium at single cell level

**DOI:** 10.1101/2023.06.01.543328

**Authors:** Pavle Vrljicak, Emma S Lucas, Maria Tryfonos, Joanne Muter, Sascha Ott, Jan J Brosens

## Abstract

Estrogen-dependent proliferation followed by progesterone-dependent differentiation of the endometrium culminates in a short implantation window. We performed single cell ATAC-seq on endometrial samples obtained across the menstrual cycle to investigate the regulation of temporal gene networks that control embryo implantation. We identified uniquely accessible chromatin regions in all major cellular constituents of the endometrium, delineated temporal patterns of coordinated chromatin remodeling in epithelial and stromal cells, and gained mechanistic insights into the emergence of a receptive state through integrated analysis of enriched transcription factor (TF) binding sites in dynamic chromatin regions, ChIP-seq analyses, and gene expression data. We demonstrate that the implantation window coincides with pervasive cooption of transposable elements (TE) into the regulatory chromatin landscape of decidualizing cells and expression of TE-derived transcripts in a spatially defined manner. Our data constitute the first comprehensive map of the chromatin changes that control TF activities in cycling endometrium at cellular resolution.

## INTRODUCTION

In humans and other simians, the uterine mucosa - endometrium - is subjected to iterative cycles of menstrual breakdown and repair.^1^ In each ovulatory cycle, menstruation is followed by rapid proliferation of resident epithelial and stromal cells in response to rising ovarian estradiol production. Proliferation in human endometrium peaks around day 10 of the cycle and leads to the formation of a superficial layer,^2^ which on average quadruples the thickness and volume of the endometrium.^3^ Following ovulation, proliferation in glandular epithelium first decreases and then ceases altogether, reflecting a switch in ovarian hormone production to progesterone. The onset of apocrine glandular secretion coincides with an abrupt change in gene expression,^4^ indicative of an epithelial cell stress response that marks the start of the mid-secretory implantation window.^5,6^ An acute stress response, termed ‘decidual reaction’, is also apparent in the stroma, characterized histologically by transient edema, angiogenesis, and influx and proliferative expansion of uterine natural killer (uNK) cells.^1,6^ In parallel, inflammatory reprogramming of stromal cells over approximately four days leads to the emergence of stress­resistant, progesterone-dependent decidual cells, heralding closure of the implantation window.^7^ Upon embryo implantation, sustained progesterone signaling enables decidual cells to engage uNK cells, resulting in pruning of stressed and senescent fibroblasts and transformation of the endometrium into the decidua of pregnancy,^7–9^ a tolerogenic matrix in which local immune cells cooperate with invading placental trophoblast to form a hemochorial placenta.^10,11^ In non-conception cycles, however, decidual cells switch to a senescent phenotype in response to falling progesterone levels, triggering sterile inflammation, influx of neutrophils and macrophages, tissue breakdown and menstrual shedding of the superficial endometrial layer.^1^

Ovarian hormones exert overall control over endometrial dynamics across the menstrual cycle through binding and activation of their cognate nuclear receptors in resident stromal and epithelial cells. The estrogen receptor 1 (ESR1) and progesterone receptor isoforms PGR-A and PGR-B (encoded by a single gene, *PGR*) are members of the nuclear receptor superfamily of transcription factors (TFs). Upon activation, the receptors dimerize, bind canonical DNA response elements in promoter or distal enhancer regions of target genes, and recruit cofactors and coregulators (i.e., coactivators and corepressors) to activate or silence gene expression.^12^ Estrogen dependent proliferation of the endometrium is spatially restricted and becomes more pronounced with increasing distance away from the basal layer, likely reflecting the presence of morphogen and cytokine gradients involved in tissue patterning and cell specification.^1^ Following ovulation progesterone-dependent differentiation is restricted to the superficial layer and mediated by complex local autocrine and paracrine signals, induction of evolutionarily conserved TFs, and sequential activation of cell-specific gene networks.^6,7^ Approximately five days after ovulation, the endometrium becomes receptive to embryo implantation. Timing of the implantation window is critical as it aligns embryonic development to the blastocyst stage with a maternal environment supportive of post-implantation development. Consequently, asynchronous embryo implantation is considered a major cause of infertility and early pregnancy loss,^13^ although the underlying pathological mechanisms remain poorly understood.

How the myriad of hormone-dependent signals and effectors are integrated to produce cell­specific and time-sensitive transcriptional responses is a major unresolved issue in endometrial biology. Endometrial differentiation coincides with altered histone modifications and other epigenetic changes,^6,14,15^ which control chromatin structure, that is, packaging of genomic DNA into nucleoprotein complexes. The organization of chromatin into sites of accessible and restricted DNA is cell-specific and highly dynamic, with structural transitions reportedly occurring on timescales that range from milliseconds to minutes and hours.^16^ Thus, the ability of TFs and cofactors to regulate gene expression is spatiotemporally constrained by accessibility to specific *cis*-regulatory elements in promoter and enhancer regions. In this study, we subjected timed endometrial biopsies to single cell Assay for Transposase-Accessible Chromatin with sequencing (scATAC-seq).^17^ We demonstrate that endometrial differentiation is underpinned by coordinated ‘waves’ of chromatin remodeling that demarcate networks of accessible sites enriched in specific TF binding sites (TFBSs). Further, dynamic changes in chromatin accessibility span the entire secretory phase of the menstrual cycle and lead to cooption of transposable elements (TEs), including Alu, LINE-1 (L1) and other elements, into the regulatory landscape of decidualizing stromal cells during the implantation window. Thus, dynamic changes in chromatin accessibility codify the multitude of signals implicated in cyclical endometrial remodeling into robust hormone-dependent gene expression programs.

## RESULTS

### Identification of endometrial cell types by scATAC-seq

Flash frozen endometrial biopsies obtained on different days of the menstrual cycle were subjected to scATAC-seq (Figure 1A). Demographic information on study participants is presented in Table S1. Proliferative phase endometrial biopsies were timed relative to the start day of menstruation (n=2, designated P7 and P10), whereas secretory phase samples were obtained on different days following the pre-ovulatory luteinizing hormone (LH) surge as determined by home ovulation testing. Each secretory phase biopsy was assigned the equivalent day of a standardized 28-day cycle (n=10, designated S18-S25). To ensure accurate timing of the post-ovulatory biopsies, transcript levels of two epithelial genes, *GPX3* and *SLC15A2*, were measured. *GPX3* and *SLC15A2* are up- and down-regulated, respectively, upon progression of the secretory phase,^18^ making the ratio of expression levels a convenient marker of ‘endometrial time’. Further, two biopsies each were obtained on cycle days 19 and 21 (S19/S19a and S21/S21a) and included as timing controls.

**Figure 1.**
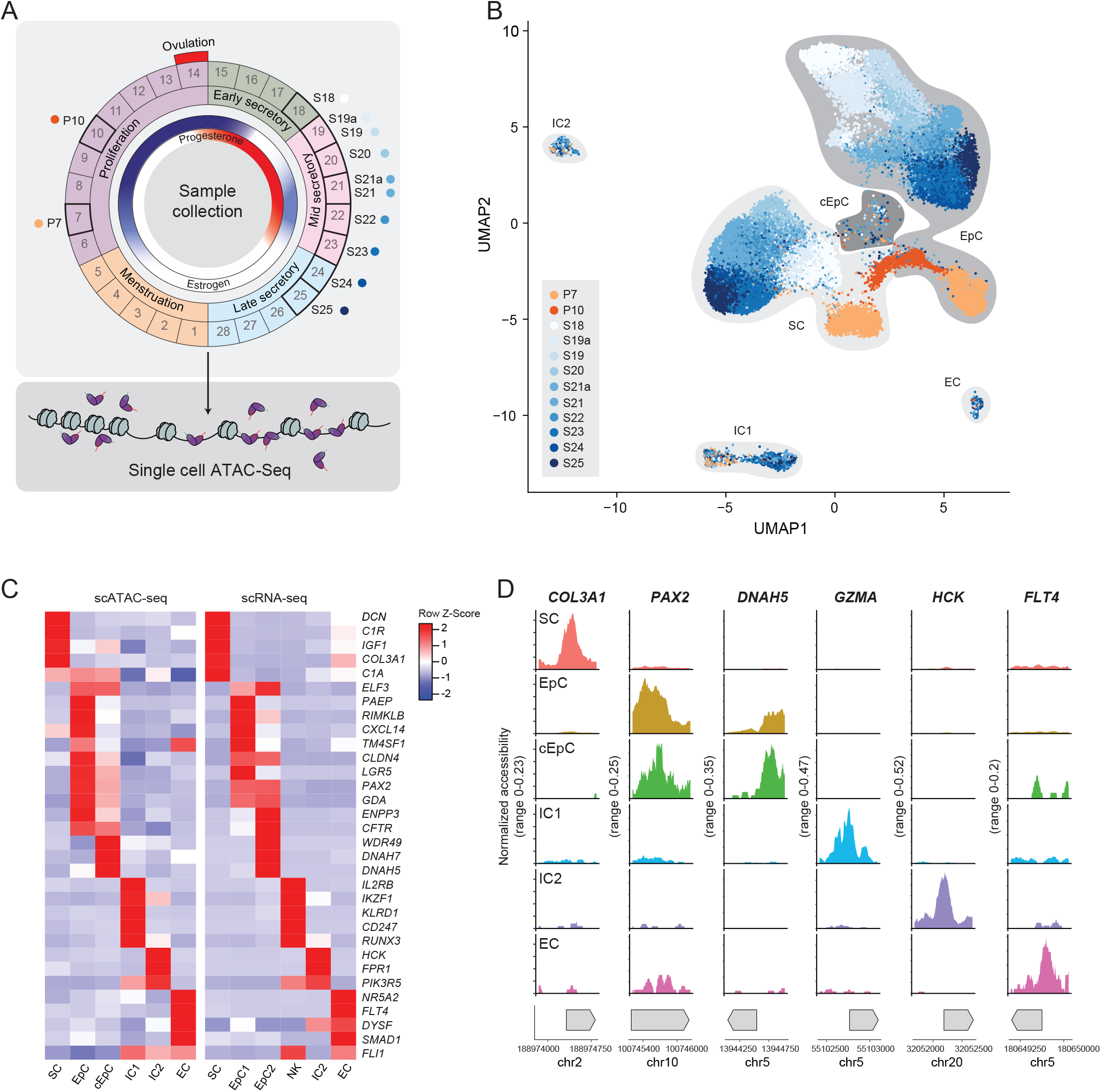
scATAC-seq analysis of cycling human endometrium. (A) Schematic illustrating the menstrual cycle, ovarian hormone levels, and day of endometrial sample collection. Proliferative samples were collected at 7 and 10 days (P7 and P10) after the start of the last menstrual period. Secretory samples were obtained 5 to 12 days following the pre-ovulatory LH surge and designated the equivocal day of a standardized 28-day cycle (S18-S25). (B) Endometrial cell clusters from ATAC-seq analysis visualized by UMAP and colored by the cycle day of samples. Cell-types were identified as stromal cells (SC), endothelial cells (EC), immune cells (IC1 and IC2) and ciliated or unciliated epithelial cells (cEpC and EpC, respectively). (C) Heatmap showing cell type-specific inferred RNA expression (z-score) by scATAC-seq and expression of corresponding transcripts in scRNA-seq data. The data on transcript levels were extracted from the study of Lucas *et al*. ^7^ (D) Examples of differential accessibility to regulatory chromatin regions at canonical marker genes in different endometrial cell types.

A total of 36,177 nuclei passed quality control. Following clustering and UMAP (uniform manifold approximation and projection) dimensionality reduction, 6 major cell clusters with unique patterns of accessible chromatin were identified (Figures 1B and 1C). Cross-referencing of inferred gene expression levels based on chromatin accessibility with known marker genes of endometrial cell types readily identified the clusters as stromal cells (SC; ∼46.6% of total nuclei), epithelial cells (EpC; ∼44.4%), ciliated epithelial cells (cEpC; ∼1.2 %), endothelial cells (EC; ∼1.2%), and two major immune cell populations (IC1; ∼4.8%, and IC2; ∼1.8%), corresponding to uNK cells and monocytes/macrophages, respectively (Figure 1D).^7^ Evidence of temporal changes in chromatin accessibility in stromal and epithelial cells was obvious from UMAP visualization, with pre- and post-ovulatory populations forming distinct clusters and the appearance of a gradient based on timing of samples across the secretory phase of the cycle. Further, the chromatin accessibility profiles of samples obtained on the same menstrual cycle day (S19/S19a and S21/S21a) aligned closely.

### Transition from the proliferative to secretory endometrium

The ovulatory shift in ovarian hormone production from estradiol to progesterone controls the transition from proliferative to secretory endometrium. A comparison of scATAC-seq profiles between proliferative and secretory endometrial biopsies identified 5,101 dynamic chromatin regions in epithelial cells (Table S2A). With 2,146 differentially accessible regions (Table S2A), the genomic response to the change in hormonal environment was less pronounced in stromal cells. Proportionally more loci closed than opened upon transition to secretory phase of the cycle in both cell types (epithelial cells: 3,989 vs. 1099; stromal cells: 1575 vs. 571, respectively). Compared to all accessible chromatin regions, dynamic loci in stromal and epithelial cells were highly enriched in putative binding sites for ESR1 and PGR, termed estrogen- and progesterone-response elements (ERE and PRE, respectively), independent of the cycle phase (Figure 2A). For example, open chromatin regions confined to the proliferative phase were enriched in both EREs and PREs. Further, chromatin regions that became accessible following ovulation were further enriched in PREs without a reciprocal loss of putative EREs. Thus, ESR1 and PGR likely play a role in regulating endometrial gene expression in both phases of the menstrual cycle, in keeping with previous studies demonstrating ligand-independent activity of these nuclear receptors.^19,20^ Next, we subjected temporally regulated chromatin regions to *de novo* motif enrichment analysis to identify TFs involved in proliferative to secretory phase transition. In epithelial cells, ATAC-seq peaks that closed following ovulation were highly enriched in high-mobility-group (HMG)/SOX TFBSs (*p* < 10^−85^), whereas binding motifs for basic helix-loop-helix (bHLH) TFs were overrepresented in opening peaks (*p* < 10^−122^) (Figure 2B). Cross-referencing with published scRNA-seq data^4^ identified several highly expressed members of the HMG/SOX and bHLH TF families with temporal expression profiles that match the switch in TFBSs in dynamic chromatin regions (Figure 2C and Table S2B). By contrast, accessible loci restricted to proliferative stromal cells harbored an abundance of Cys2‒His2 zinc finger (C2H2-ZF) TFBS (*p* < 10^−144^) (Figure 2B). Repression of these sites following ovulation, and loss of corresponding TFs (Figure 2C), coincided with gain of binding sites for the transcriptional regulator RBPJ (*p* < 10^−34^), which functions as transcriptional repressor in the absence of Notch signaling and an activator when bound to Notch proteins.^21^ Interestingly, 262 genomic regions closed following ovulation in both epithelial and stromal cells (Table S2A). These regions were enriched in RUNT/SMAD TFBSs (*p* < 10^−12^) (Figure 2B and 2C). Whole tissue expression analysis of the nearest gene to each closing peak showed a biphasic response (Figure 2D), characterized by rapid downregulation in early-secretory endometrium followed by loss of repression prior to menstruation, reflecting the rise and fall in circulating progesterone levels, respectively. Further, gene ontology analysis revealed that genes silenced following ovulation in both the stromal and epithelial compartments are implicated in various signal transduction pathways (WNT, TGF-β, MAPK, and steroid hormone signaling) and key biological processes (e.g., growth, metabolism, and circadian gene regulation) (Figure 2E).

**Figure 2.**
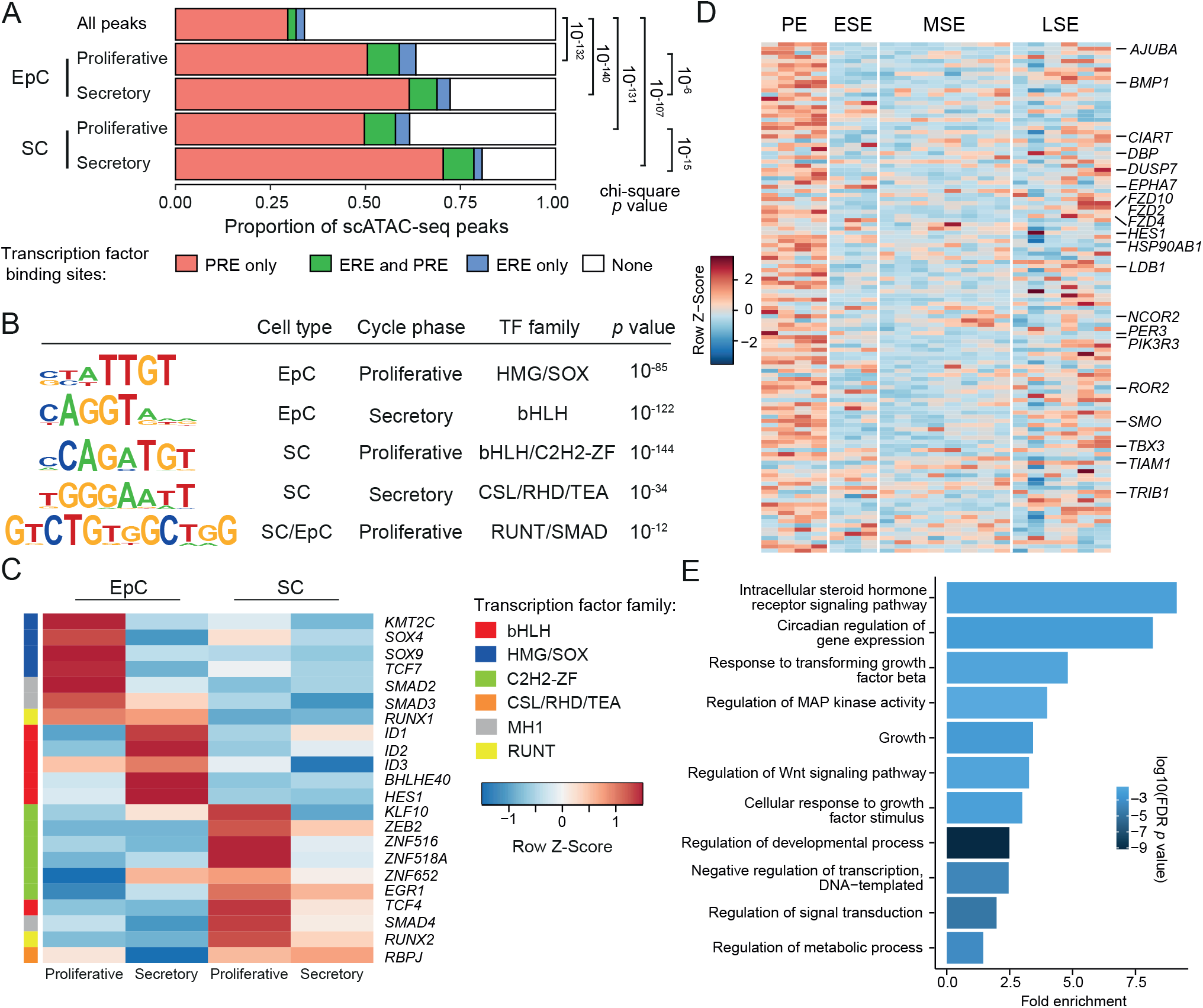
Chromatin accessibility in stromal and epithelial cells during the proliferative and secretory phase of the cycle. (A) Enrichment of ESR and PGR binding sites (ERE and PRE, respectively) in proliferative and secretory epithelial and stromal cells (EpC and SC, respectively). “All peaks” refers to every ATAC-seq peak in our dataset. (B) TF binding motifs enriched in opening and closing chromatin regions in proliferative versus secretory phase endometrium. *P* values represent binomial test results using HOMER.^65^ (C) Heatmap showing relative RNA expression (z-score) of candidate TF in proliferative and secretory phase samples. The data were extracted from the study of Wang *et al*.^4^ (D) Heatmap showing relative RNA expression (z-score) of nearest gene to each closing chromatin region in bulk endometrial gene expression data. The data were extracted from the study of Talbi *et al*.^22^ PE, proliferative endometrium; ESE, early-secretory endometrium; MSE, mid-secretory endometrium; LSE, late-secretory endometrium. (E) Gene Ontology analysis of genes silenced upon ovulation in both stromal and epithelial cells.

### Dynamic chromatin changes in differentiating epithelial cells

Next, we set out to map the temporal changes in chromatin accessibility across the implantation window in epithelial cells. The endometrium becomes receptive 5 days after ovulation for approximately 4 days (S19-S22). First, we performed pairwise comparisons of scATAC-seq peaks in epithelial cells from samples obtained between S18 and S25 to identify dynamic chromatin loci. The resulting 8,217 chromatin regions were then grouped in three temporal patterns using k-means clustering (Figure 3A and Table S3A). Pattern 1 represents 4,624 dynamic loci that close during the implantation window in a coordinated manner. Pattern 2 (n=1,645) and 3 (n=1,948) capture chromatin regions that open during the implantation window, and either remain stable or open further upon transition to the late-secretory phase of the cycle, respectively. Cross-referencing with bulk endometrial gene expression data spanning the menstrual cycle^22^ revealed that epithelial genes near closing chromatin loci in pattern 1 are downregulated in mid-secretory endometrium whereas comparable genes in patterns 2 and 3 are induced (Figure S1). While pattern 2 genes tend to peak during the mid-secretory implantation window, the expression of pattern 3 genes is maintained or enhanced upon transition to the late-secretory phase (Figure S1).

**Figure 3.**
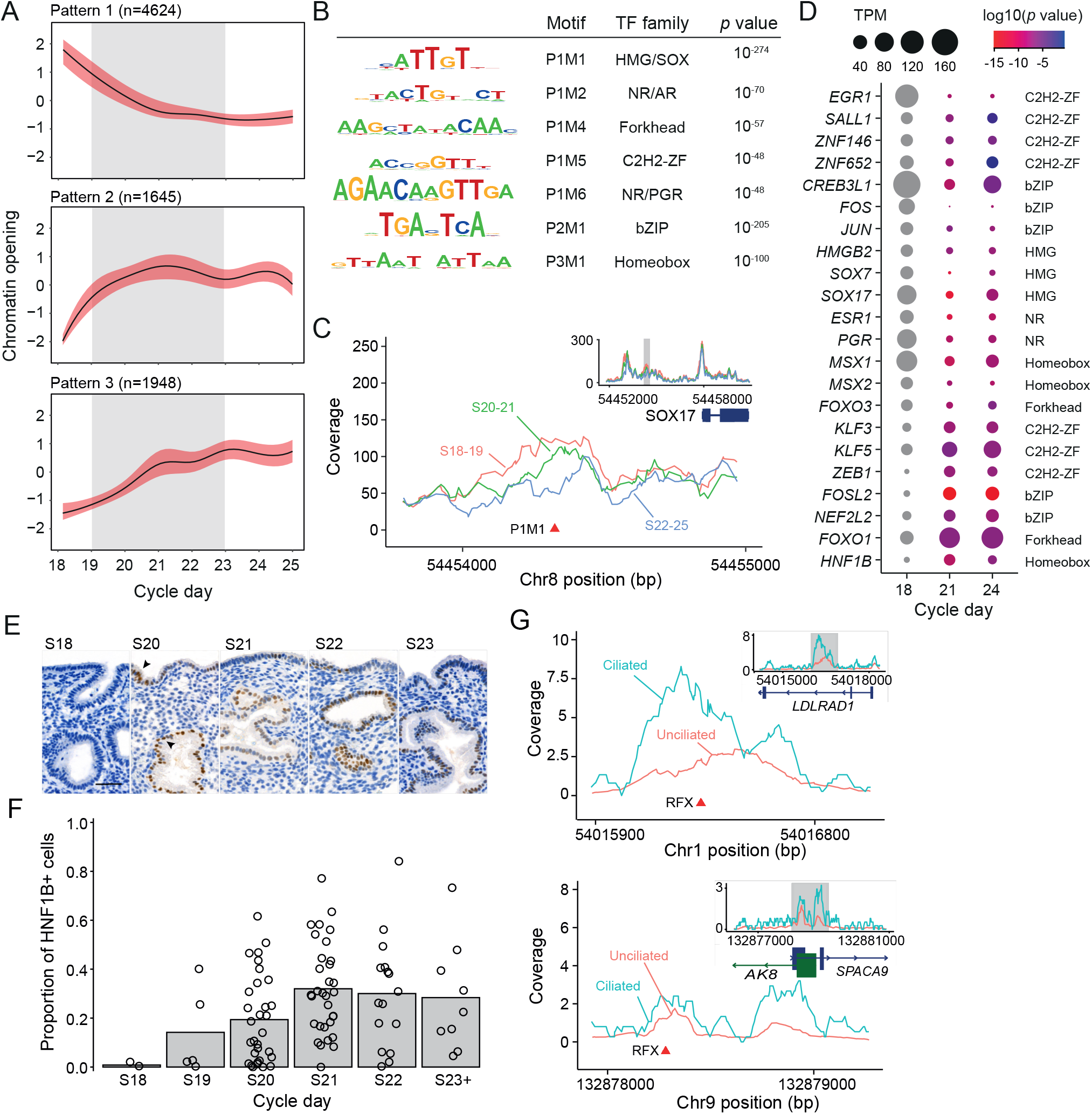
Dynamic chromatin remodeling in secretory phase epithelial cells. (A) Clustering of dynamic chromatin loci in secretory phase epithelial cells into temporal patterns. Trend curve represents LOESS fit ± standard error (red). The grey shaded area demarcates the putative implantation window. (B) TF binding motifs enriched in temporally regulated genomic regions from panel A (designated as P1M1 for <u>P</u>attern <u>1 M</u>otif <u>1</u>, etc.). *P* values represent binomial test results using HOMER.^65^ Only motifs validated by footprint analysis and enrichment over all ATAC-seq genomic regions are shown. (C) Cycle-dependent changes in chromatin accessibility upstream of *SOX17*. The location of an HMG/SOX binding site (P1M1) is indicated. (D) Expression of candidate TFs before (S18), during (S21), and after (S24) the implantation window. The data were obtained from RNA-seq analysis of laser-captured endometrial glands. ^66^ The data denote transcripts per million (TPM; Table S3) and *p* values are based on expression on day 21 (S21) and day 24 (S24) versus day 18 (S18) (Wald test using DESeq2). (E) HNF1B immunoreactivity in secretory phase endometrium. Arrows show nuclear HNF1B staining (brown). The scale bar denotes 100 µm. (F) Proportion of HNF1B-positive cells in endometrial glands across the peri-implantation window. (G) Examples of differential chromatin accessibility in ciliated versus unciliated epithelial cells. Locations of RFX binding motifs are shown.

Next, we subjected the dynamic chromatin loci in patterns 1-3 to *de novo* motif enrichment analysis, repeating the analysis 5 times with shuffled chromatin regions to establish a robust threshold for statistical significance for each pattern. A total of 11 significant motifs were identified, eight in pattern 1 (designated P1M1-8), one in pattern 2 (P2M1), and two in pattern 3 (P3M1-2) (Figure S2A). Based on evidence of dynamic TF binding in footprint analysis (Figure S2B) and enrichment over all ATAC-seq genomic regions (Figures S2C and S2D), 7 of the 11 overrepresented motifs were deemed *bona fide cis*-regulatory elements involved in coordinated chromatin remodeling in epithelial cells across the implantation window (Figure 3B). Chromatin regions that close during the implantation window (pattern 1) were highly enriched for HMG/SOX TFBSs (*p* < 10^−274^, Figure 3B), continuing the trend observed upon transition from proliferative to secretory phase of the cycle (Figure 2B). Other closing loci in pattern 1 harbored TFBSs for Forkhead and C2H2-ZF TFs (*p* < 10^−57^ and *p* < 10^−48^, respectively) and nuclear receptors (NRs), including the PGR (*p* < 10^−48^, Figure 3B). Among the epithelial chromatin regions that open during the implantation window, pattern 2 loci were highly enriched for bZIP TFBSs (*p* < 10^−205^), whereas homeobox TF binding motifs were overrepresented in pattern 3 (*p* < 10^−100^). We noted that several cycle-dependent endometrial TFs possess dynamic TF binding motifs in their regulatory regions (Table S3B). For example, the closing chromatin locus upstream of *SOX17*, an epithelial cell enriched TF, contains an HMG/SOX binding motif (Figure 3C), thus implicating SOX17 in its own regulation. Next, we compiled a set of TFs that potentially engage enriched TF binding motifs in pattern 1-3 based on their level and temporal pattern of expression in RNA-seq data of laser-dissected endometrial glands obtained before (S18), during (S21), and after (S24) the implantation window (Figure 3D and Table S3C) (GEO ID: GSE84169). For example, *HNF1B* is the highest expressed homeobox TF gene in glandular epithelium with a temporal expression profile that follows pattern 3 closely. Immunohistochemistry of timed endometrial biopsies confirmed that increased HNF1B expression in glandular epithelium during the window of implantation is maintained upon transition to the late-secretory phase (Figures 3E and 3F).

Ciliated cells made up approximately 2.7% of the captured epithelial cell fraction in our ATAC-seq data set. While the chromatin landscape of ciliated epithelial cells is distinct, it is not subjected to obvious temporal changes across the menstrual cycle (Figure 1B). When compared to their unciliated counterparts, accessible chromatin regions in ciliated cells were enriched in 5 TF binding motifs, most prominently for the regulatory factor X (RFX) family of TFs (*P* < 10^−58^, Figure S4A). Two family members, RFX2 and RFX3, are direct regulators of core cileogenic genes and expressed selectively in endometrial ciliated epithelial cells.^23^ Figure 3G shows examples of RFX binding sites in proximity of genes (*LDLRAD1*, *AK8* and *SPACA9*) involved in cilia assembly.^24^

### Dynamic chromatin changes in decidualizing stromal cells

The mid-secretory implantation window coincides with decidualization, a process defined by inflammatory reprogramming of stromal cells into specialized decidual cells. The emergence of phenotypic decidual cells on day 23 (S23) of a standardized 28-day cycle, characterized by their rounded appearance and enlarged nuclei, marks the closure of the window.^5^ Using the same approach as for epithelial cells, we resolved the 2,392 dynamic chromatin loci in stromal cells between S18 and S25 into 4 temporal patterns (Patterns 1-4, Figure 4A and Table S4). Chromatin dynamics in decidualizing stromal cells were notably more variable when compared to differentiating epithelial cells, likely reflecting that endometrial glands are clonal in origin and therefore subject to tighter regulatory control.^25^ Opening of the implantation window was marked by a sequential shift in peak accessibility from pattern 1 to pattern 2 and 3, whereas closure of the window coincided with maximal accessibility in pattern 4. Using *de novo* motif analysis, we identified 16 significantly overrepresented TF motifs across the four patterns (Figure S3A), 12 of which were considered primary *cis*-regulatory elements based on footprint analysis (Figure S3B) and enrichment over all ATAC-seq genomic regions (Figure S3C and 3D). As shown in Figure 4B, patterns 1 and 4, which mark the boundaries of the implantation window, were selectively enriched in C2H2-ZF and bZIP TFBSs, respectively. Multiple motifs were overrepresented in patterns 2 and 3, likely reflecting active reprogramming of stromal cells into pre-decidual cells. Next we mined scRNA-seq data to identify TFs in stromal cells that are temporally regulated across the peri-implantation window.^4^ In keeping with our scATAC-seq data, several genes encoding for C2H2-ZF TFs of the Krüppel-like factor (KLF) family (pattern 1) are downregulated upon opening of the implantation window, which is followed by induction of genes encoding known core TFs in pre-decidual cells, including HOXA (Homeobox A Cluster, pattern 2) and STAT (signal transducer and activator of transcription) family members (pattern 3) (Figure 4C). Notably, pattern 2 is also enriched in NR binding sites. Apart from estradiol and progesterone, decidualizing stromal cells are responsive to androgen and corticosteroid signaling, at least in primary cultures.^19,26^ In line with these observations, all steroid hormone receptor genes, including *ESR1*, *PGR*, *AR* (androgen receptor), *NR3C1* (glucocorticoid receptor), and *NR3C2* (mineralocorticoid receptor), are dynamically expressed in pre-decidual cells across the implantation window *in vivo* (Figure 4C). Thus, a notable level of congruency was observed between accessibility to TFBSs in patterns 1-3 and the temporal expression of corresponding TFs. However, congruency was less obvious for pattern 4, which captures the dynamic chromatin regions in emerging decidual cells around day 23 of a standardized 28-day cycle.

**Figure 4.**
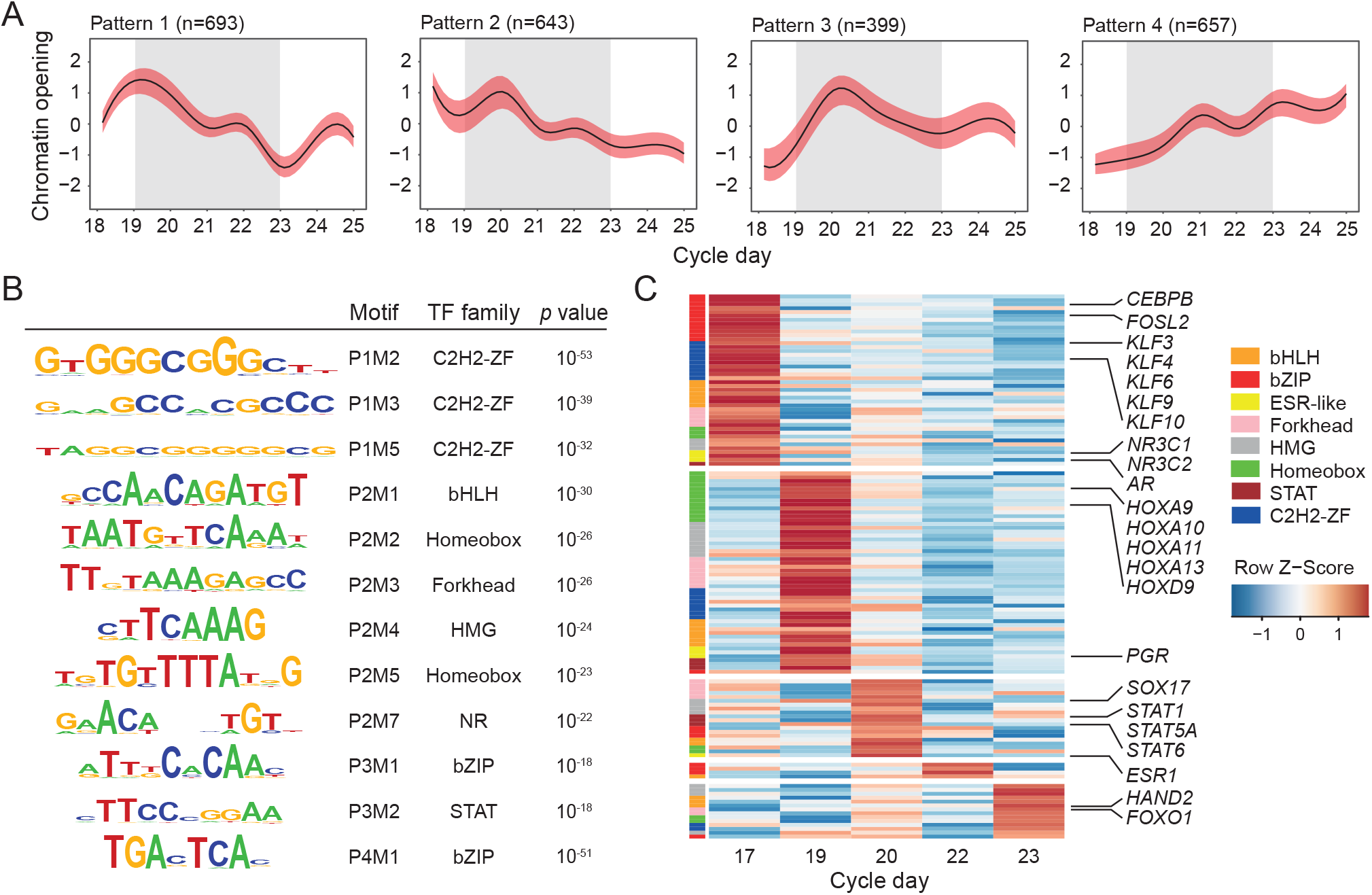
Dynamic chromatin remodeling in secretory phase stromal cells. (A) Clustering of dynamic chromatin loci in secretory phase stromal cells into temporal patterns. Trend curve represents LOESS fit ± standard error (red). The grey shaded area demarcates the putative implantation window. (B) TF binding motifs enriched in temporally regulated genomic regions from panel A (designated as P1M1 for <u>P</u>attern <u>1 M</u>otif <u>1</u>, etc.). *P* values represent binomial test results using HOMER.^65^ Only motifs validated by footprint analysis and enrichment over all ATAC-seq genomic regions are shown. (C) Relative expression (z-score) of candidate TFs. The data were extracted from the study of Wang *et al*.^4^ See also Table S4.

### Cooption of transposable elements during the implantation window

Novel *cis*-regulatory elements encoded by Mammalian-, Therian-, and Eutherian-specific TEs played an important role in the emergence of decidual cells in ancestral eutherian (placental) mammals.^27,28^ A second wave of TE cooption in Catarrhine primates (Old World monkeys, apes, humans) has been linked to the emergence of novel reproductive traits, such as spontaneous decidualization of the endometrium (i.e., independently of embryo implantation) and menstruation.^29^ To gain insights in the regulation of decidualizing stromal cells, we mapped enriched TEs across the temporal scATAC-seq peaks (patterns 1-4). Progressive differentiation of stromal cells into pre-decidual (pattern 3) and decidual cells (pattern 4) coincided with repression of low-complexity elements and simple repeats and enrichment of distinct TE families across the 5 major classes: DNA transposons, Short Interspersed Nuclear Elements (SINE), Long Interspersed Nuclear Elements (LINE), composite SVA (SINE-VNTR-Alu) elements, and long terminal repeats (LTR) retrotransposons (Figure 5A and Table S5). Intriguingly, when compared to pre-decidual cells (pattern 3), accessible chromatin loci in decidual cells (pattern 4) are characterized by overdispersion of evolutionarily young TE families and subfamilies (e.g., SVA, L1PA, and AluY) and loss of some older elements (e.g., MIR and hAT.Charlie) (Figure 5A). Perhaps the most salient feature of decidualizing stromal cells is the cooption of primate specific Alu elements, which make up ∼10% of the human genome. Compared to ATAC-seq peaks in stromal cells (pattern 1), enrichment in decidual cells (pattern 4) was 1.53-fold for AluJ, the oldest of the Alu lineages, 1.52-fold for AluS, and 3.15-fold for AluY, the youngest lineage which expanded in Catarrhine primates.^30^ Alu elements are 300 bp in length and composed of a left and right monomer on either side of an A-rich linker region, and a 3’ poly-A tail. The A and B boxes in the left monomer contain RNA polymerase III promoter binding sites. In decidualizing stromal cells, accessibility to Alu elements increases in a time-dependent manner (Figure 5B). Integrative analysis of putative *cis*-regulatory elements and ChIP-seq data obtained from primary endometrial stromal cell cultures^31–34^ and the ENCODE database^35^ confirmed binding of decidual TFs, including GATA2, FOXO1 and CEBPB,^6,36^ to a cluster of canonical TFBSs in the right monomer of opening *Alu* elements (Figure 5B). ChIP-seq data also demonstrated prominent binding of ESR1, although not PGR, in line with TFBS predictions (Figure 5B and Figure S5).

**Figure 5.**
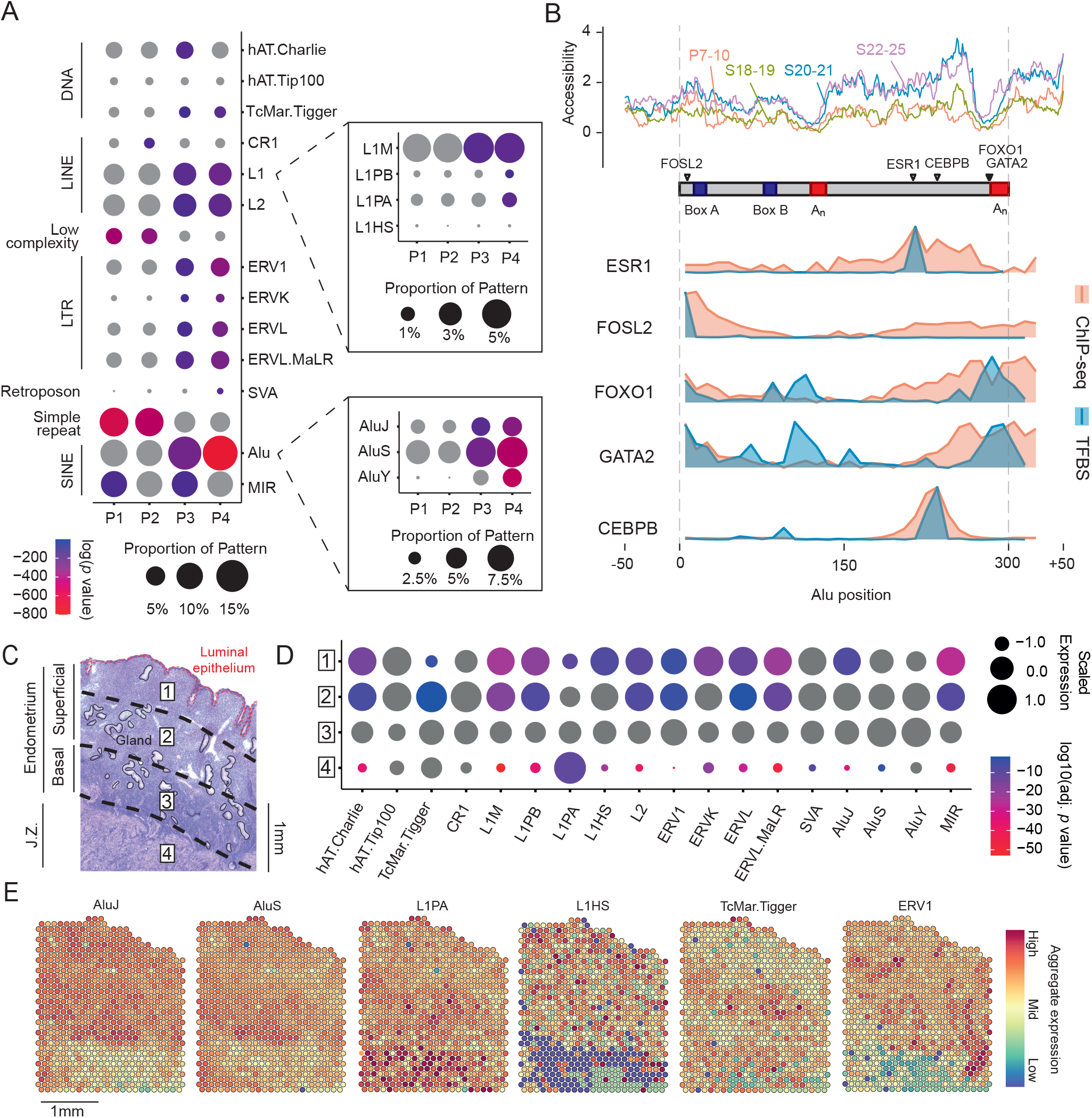
Transposable elements in dynamic chromatin regions in decidualizing stromal cells. (A) Enrichment of transposable elements (TEs) in the temporal chromatin waves in decidualizing stromal cells. The size of the circles denotes the proportion of TEs, and the color key indicates *p* values (hypergeometric test for enrichment). (B) Analysis of Alu element showing boxes A and B (blue) and A-rich regions (red). Cycle­dependent changes in chromatin accessibility are color coded and labelled by the cycle day of the sample. Density plots showing location of predicted and validated TFBSs from JASPAR database and ChIP-seq peak summits, respectively. (C-E) Spatial transcriptomic analysis of an early-secretory phase full-thickness endometrial sample with adjacent junctional zone myometrium.^37^ (C) Hematoxylin and eosin (H&E) staining of the tissue sample. For transcriptomic analysis, the sample was divided into 4 regions, broadly corresponding to the superficial endometrial layer (regions 1 and 2), the regenerative basal layer (region 3), and junctional zone myometrium (region 4). (D) Normalized average expression of TE-derived transcripts across the four regions. The size of the circles corresponds to the change in relative expression from the mean. The color key denotes the adjusted *p* value in one region against the other three regions (Wilcoxon rank sum test). (E) Examples of spatial expression of TE-derived transcripts across the endometrium and junctional zone myometrium.

Endometrial responses to steroid hormone signaling are spatially organized and largely confined to the superficial endometrium.^1^ Hence, we examined the expression of TE-derived transcripts in spatial transcriptomic data (Visium) obtained from a surgical specimen containing full-thickness early-secretory endometrium and adjacent myometrium, also termed ‘junctional zone’ myometrium.^37^ We used a modified pipeline to count TE-derived reads in a locus-specific manner^38^ across four zonal uterine regions (Figure 5C). Regions 1 and 2 represent the superficial endometrial layer whereas regions 3 and 4 capture broadly the basal endometrial layer and junctional zone myometrium, respectively. As shown in Figures 5D and 5E, TE-derived transcripts, with exception of L1PA, are more abundantly expressed in the endometrium than junctional zone myometrium. However, L1HS (previously known as L1PA1), the evolutionarily youngest and the only active L1 subfamily in the human genome,^39^ is expressed most prominently in subluminal endometrium. Further, even within the superficial endometrium (regions 1 and 2), there was evidence of a gradient in the expression of TE-derived transcripts (Figures 5D and 5E). To substantiate these observations, we measured the expression of L1 mRNA in 36 RNA-seq libraries derived from endometrial biopsies obtained between days 18 and 25 of the cycle.^18^ L1 RNA is bicistronic, encoding two non-overlapping open reading frames, ORF1 and ORF2, whose protein products (ORF1p and ORF2p) bind L1 RNA to form a ribonucleoprotein complex.^40^ Interestingly, L1 ORF transcript levels correlated closely with principal component 3 (PC3), which accounts for 6.64 % of variance in gene expression in this set of 36 endometrial samples (Figure S6). Unlike PC1 and PC2, which capture temporal gene expression changes in cycling endometrium,^18^ PC3 reflects spatial distribution of transcripts across the luminal-basal axis.

### Endometrial endothelial and immune cell populations

The endometrium is one of few adult tissues that undergo cycles of physiological angiogenesis.^41^ Although endometrial endothelial cells express ESR1 and PGR,^42^ we did not observe a compelling temporal scATAC-seq signature (Figure 6A). However, vascular and lymphatic endothelial cell clusters were readily identified based on differences in chromatin accessibility (EC1 and EC2, respectively; Figure 6A). When compared to the lymphatic ATAC-seq peaks, accessible chromatin regions in vascular endothelial cells were selectively enriched in multiple motifs, including bZIP (*p* < 10^−81^), Homeobox (*p* < 10^−75^), and interferon-regulatory factor (IRF; *p* < 10^−68^) TFBSs (Figure 6B).

**Figure 6.**
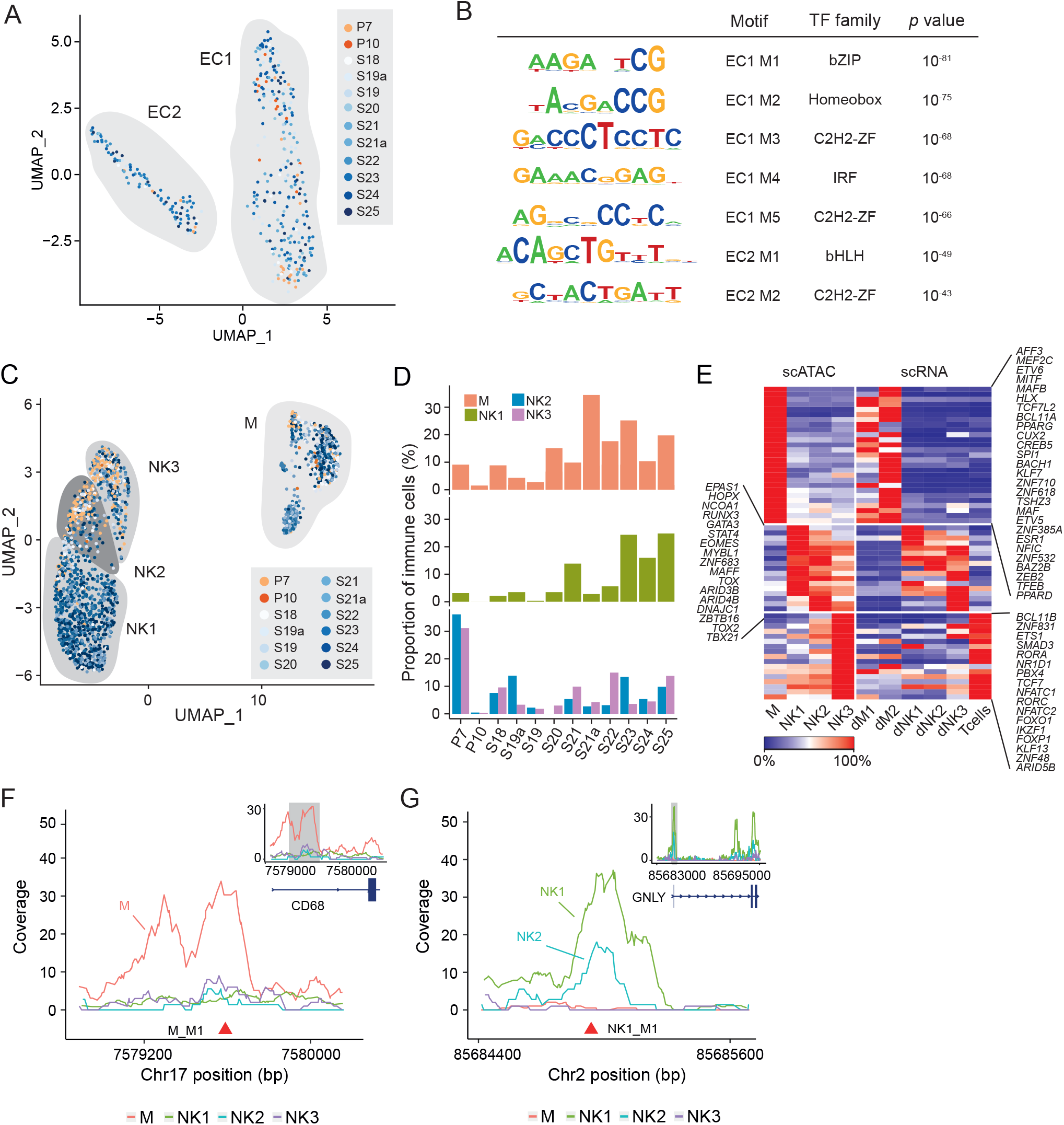
Chromatin accessibility in endometrial endothelial and immune cells. (A) UMAP visualization of vascular and lymphatic endothelial cells (EC1 and EC2, respectively), color coded by the day of the sample in the menstrual cycle. (B) TF binding motifs enriched in differentially accessible chromatin regions of EC1 and EC2 nuclei. (C) UMAP visualization of endometrial macrophages (M) and natural killer cells (NK) color coded by the cycle day of the samples. (D) Heatmap showing relative inferred RNA expression (z-score) of candidate cell-type specific TFs and expression of corresponding transcripts in scRNA-seq data. Designations of cell populations and transcript levels were extracted from the study of Vento-Tormo *et al*. ^11^ (E) Change in immune cell subtype proportions across the secretory phase of the menstrual cycle. (F) Coverage plot showing differential chromatin peaks in endometrial macrophages and NK subsets. Regulatory regions of *CD68* and *GNLY* are shown with location of corresponding enriched TFBSs.

Uterine immune cells, especially uNK cells, play a pivotal role in endometrial remodeling before and after embryo implantation.^8,11^ We identified 3 subsets of uNK cells, termed NK1-3, based on differences in chromatin accessibility (Figure 6C). The NK2 and NK3 subsets were prominent in the post-menstrual endometrial sample (P7), but their relative abundance appeared stable across the remainder of the cycle. By contrast, macrophages and NK1 cells were more abundant in mid- and late-secretory endometrium (Figure 6D). Cross-referencing of top inferred differentially expressed genes with transcriptomic profiles obtained from immune cells at the maternal-fetal interface in early pregnancy^11^ confirmed that NK1 cells correspond to cytokine-producing decidual NK1 (dNK1) cells (Figure 6E). This NK subset is characterized phenotypically by cell surface expression of killer cell immunoglobulin-like receptors (KIRs) and the ectonucleotidase CD39 (ENTPD1).^43^ NK2 cells correspond to the intermediate dNK2 subset in early pregnancy, whereas cytotoxic NK3 cells are akin to dNK3 and T cells (Figure 6E). Next, we integrated *de novo* motif enrichment analysis and TF expression profiles to gain insight into the transcriptional regulation of endometrial immune cells. We identified 5 TF binding motifs selectively enriched in macrophages when compared to all uNK subsets (Figure S4B). Figure 6F shows the location of the most enriched motif (ETS, designated M-M1; *p <* 10^−95^) in macrophages in the proximal promoter region of *CD68*, the canonical marker of human monocytes and tissue macrophages. We also identified 5 motifs selectively enriched in NK1 cells when compared to NK2 or NK3 subsets (Figure S4B). Figure 6G shows the location of the most enriched motif, a RUNT binding site designated NK1-M1 (*p <* 10^−64^), upstream of the transcriptional start site of *GNLY*, encoding granulysin. In keeping with the level of accessibility at this locus, *GNLY* transcript levels are high, intermediate and low in NK1, NK2 and NK3 cells, respectively.^11^

## DISCUSSION

We demonstrated that all cell types in the endometrium have a unique genomic organization that confers access to cell-specific combinations of enriched TFBSs. We also found that the chromatin landscapes of resident epithelial and stromal cells are exquisitely responsive to changes in circulating ovarian hormone levels, exemplified by the observation that many more genomic regions close than open upon transition from the proliferative to the secretory phase of the cycle. We identified a set of 262 common loci, enriched in RUNT/SMAD binding motifs, which close simultaneously in stromal and epithelial cells following ovulation. Gene ontology analysis of the nearest genes revealed overrepresentation of modulators of multiple signal transduction pathways, including WNT and steroid hormone signaling, and core biological processes, such as metabolism, growth, and circadian gene regulation. Despite the overall reduction in open chromatin following ovulation, access to PREs was conspicuously enriched in resident endometrial cells. Further, the emergence of a receptive endometrial state during the implantation window was underpinned by highly coordinated changes in chromatin accessibility. In epithelial cells, we discerned two distinct waves of opening chromatin, encompassing 1,645 and 1,948 loci, respectively, concurrent with loss of accessibility at 4,624 genomic sites. Similarly, decidual reprogramming of stromal cells during the implantation window involved closure of 693 genomic regions and three distinct waves of opening chromatin, each comprising hundreds of loci and peaking at different timepoints. The waves of opening and closing chromatin in both cell types resulted in time-sensitive enrichment and depletion, respectively, of active TFBSs as demonstrated by footprint analyses. Put differently, our findings revealed that the implantation window coincides with a coordinated switch in the regulatory DNA landscape of resident endometrial cells. To gain further insights, we integrated our scATAC-seq findings with cell-specific changes in endometrial TF expression. Although this exercise does not prove that a given TF is active at a specific site, we nevertheless observed a notable degree of congruency between the gain or loss of specific *cis*-regulatory elements in dynamic chromatin regions and expression of corresponding TF family members. For example, decidual reprogramming of stromal cells during the implantation window was associated with sequential loss of C2H2-ZF motifs and enrichment of binding sites for - amongst others - Homeobox and STAT TFs, which coincided with repression of Krüppel-like TFs and induction of known decidual transcriptional regulators, including multiple STAT and HOXA family members.^44–46^ In epithelial cells, we identified FOSL2, an important PGR coregulator,^33^ and HNF1B as putative drivers of glandular differentiation based on the alignment between expression of these TFs and enrichment of corresponding regulatory elements in opening chromatin regions.

Rapid evolution of reproductive processes in eutherians (placental mammals) is attributed to exaptation of species-specific TEs into the regulatory DNA landscape of pre-implantation embryos, placental cell lineages and maternal decidual cells,^29,47^ resulting in the gain and loss of entire gene sets between species.^48^ While decidual cells likely originated in ancestral eutherians in parallel with invasive hemochorial placentation,^49^ cooption of novel TE-derived *cis*-regulatory elements in Catarrhine primates has been linked to the emergence of specific endometrial traits, including heightened progesterone responsiveness, spontaneous decidualization and menstruation, and deep hemochorial placentation.^29^ In accordance, we observed pervasive enrichment of TEs in opening chromatin regions upon differentiation of stromal cells into pre-decidual and decidual cells, a process that marks the boundaries of the implantation window. Further, evolutionarily ancient and young TEs were coopted sequentially, indicating that decidual reprogramming of human endometrial stromal cells partly recapitulates the phylogenetic roots of this differentiation process. For example, increased accessibility to Alu elements upon differentiation of pre-decidual into decidual cells favored relative enrichment of young (AluY) over older elements (AluS and AluJ). In line with previous observations in primary cultures,^50^ exaptation of Alu elements in the regulatory DNA landscape of decidual cells enables evolutionarily core TFs, including CEBPB, FOXO1 and GATA2,^51^ to access a composite response element comprising multiple TFBSs near the 3’ end of the right monomer. This observation provides further evidence that decidual gene expression is under control of multimeric TF complexes,^6,52,53^ which in the case of Alu involves DNA binding of ESR1 rather than PGR.

We also demonstrated that accumulation of TE-derived transcripts is a physiological phenomenon in cycling human endometrium, especially in the decidualizing superficial layer. Retrotransposons are firmly repressed in most adult tissues because of the deleterious consequences associated with reactivation, including retrotransposition of L1 or Alu elements, genomic instability and DNA damage, transcriptional interference, and innate immune activation in response to cytoplasmic accumulation of free DNA or dsRNA.^54,55^ However, de­repression of evolutionarily young retrotransposons and increased transcription are also hallmarks of replicative cellular senescence.^56,57^ Interestingly, spontaneous decidualization of endometrial stromal cells not only follows intense proliferation prior to ovulation but starts with an acute cellular stress response, characterized by a burst of reactive oxygen production (ROS) and release of inflammatory mediators.^7,58,59^ Further, decidual cells share multiple characteristics with canonical senescent cells, including cell cycle arrest, expression of survival genes, heightened autonomy from environmental cues, and rounded appearance with abundant cytoplasm and enlarged nuclei.^6^ A crucial difference, however, is the suppression of a pro-inflammatory senescence-associated secretory phenotype (SASP) in decidual cells, likely reflecting progesterone-dependent silencing of key pro-inflammatory pathways, including MAPK/JNK,^60^ NF-κB,^61^ and type 1 interferon signaling.^62,63^ Thus, rather than being a novel cell type that originated in ancestral eutherians, as proposed by others,^51,64^ decidual cells may represent endometrial stromal cells in which progesterone exerts control over the senescence phenotype and firmly suppresses SASP activation. In keeping with this conjecture, falling progesterone levels prior to menstruation triggers a rapid phenotypic switch in decidual cells, characterized by a rise in P16^INK4a^-positive cells, secretion of extracellular matrix proteinases, inflammatory mediators, and chemokines, and recruitment of macrophages and neutrophils, which in turn reinforce cellular senescence through ROS production.^1^ Whether retrotransposition occurs at higher frequency in the endometrium than in other somatic tissues warrants further exploration. However, it is notable that intrinsic mechanisms operate in the endometrium that *a priori* mitigate against the deleterious impacts of transposon activity, including an abundance of uNK cells involved in immune surveillance and clearance of stressed and damaged stromal cells and the wholesale disposal of the decidualizing endometrium at menstruation or following parturition.^1^

### Limitations of the study

Although we carefully timed the endometrial biopsies for scATAC-seq analysis, the sample size was limited and insufficient to assess physiological variability. The biopsies, which contain only superficial endometrium, also did not cover the menstrual phase of the cycle. Further studies are needed to map the changes in DNA methylation and histone modifications associated with dynamic chromatin remodeling and to parse the underlying regulatory mechanisms. We were only able to infer the impacts of enriched binding motifs in opening and closing scATAC-seq peaks on TF activity and gene expression. Additional genomic investigations, including mapping of cell type-specific enhancers, are needed to gain granular insights into how cell-specific chromatin changes control the spatiotemporal gene expression in the endometrium. Although our observation of widespread TE de-repression and transcription signals a novel dimension in endometrial physiology, the impact of this phenomenon on reproductive health and disease warrants further investigations.

In conclusion, by mapping chromatin accessibility at single cell level, we demonstrated that cyclical endometrial remodeling involves highly coordinated changes in chromatin accessibility in endometrial cells, which converge during the implantation window. In stromal cells, this convergence constitutes an inflection point, away from sequential enrichment of specific DNA binding motifs and expression of corresponding TFs and towards cooption of TE-derived complex response elements evolved to assume control over decidual gene expression. Our data constitute a new resource for the exploration of gene regulation in cycling human endometrium and the benchmarking of mechanistic studies *in vitro*.

## ACKNOWLEDGMENTS

We wish to thank patients attending the Implantation Research Clinic at University Hospitals Coventry and Warwickshire (UHCW) National Health Service Trust (NHS) for contributing endometrial samples for research. We also thank University of Birmingham and University of Warwick Genomics facilities and the staff of the Biomedical Research Unit at UHCW NHS Trust. We are indebted to Dr Katherine Fishwick for assistance with immunohistochemistry. This work was supported by funds from the Tommy’s National Miscarriage Research Centre and a joint Wellcome Trust Investigator Award to J.J.B. and S.O. (212233/Z/18/Z).

## Author contributions

Conceptualization: J.J.B. and S.O.; Software: P.V.; Methodology: P.V. and E.S.L.; Formal Analysis: P.V. and J.J.B.; Investigation: P.V., J.J.B., E.S.L., J.M., and M.T.; Resources: J.J.B.; Data curation: P.V., J.J.B., E.S.L and J.M.; Writing ‒ Original Draft: J.J.B. and P.V.; Writing ‒ Review and Editing: J.J.B., P.V., E.S.L. and J.M.; Visualization: P.V., E.S.L. and J.M. Supervision: J.J.B. and S.O. Funding Acquisition: J.J.B. and S.O.

## Declaration of interests

The authors declare no competing interests.

## SUPPLEMENTAL INFORMATION

**Table S1.** Demographic details of tissue donors.

**Table S2.** Chromatin changes in epithelial and stromal cells upon transition from proliferative to secretory phase of the menstrual cycle.

(A) Dynamic ATAC-seq peaks in proliferative versus secretory samples and associated gene regions.

(B) Transcription factors expressed differentially in proliferative and secretory phase epithelial and stromal cells.

**Table S3.** Dynamic chromatin changes in differentiating epithelial cells

(A) Temporally regulated chromatin regions in secretory phase epithelial cells and their closest gene regions.

(B) Epithelial TFs containing dynamic binding motifs in their regulatory regions. RNA-seq expression of closest genes in endometrial glands before (S18), during (S21) and after (S24) the implantation window (GEO ID: GSE84169).

(C) Temporal expression of TFs in endometrial secretory phase glands.

**Table S4.** Dynamic chromatin changes in decidualizing stromal cells.

(A) Temporally regulated chromatin regions in secretory phase stromal cells and their closest protein coding regions.

(B) Temporal expression of TFs in secretory phase stromal cells (data for Figure 4C).

**Table S5.** Repetitive element enrichment in stromal cell temporal patterns.

**Figure S1.**
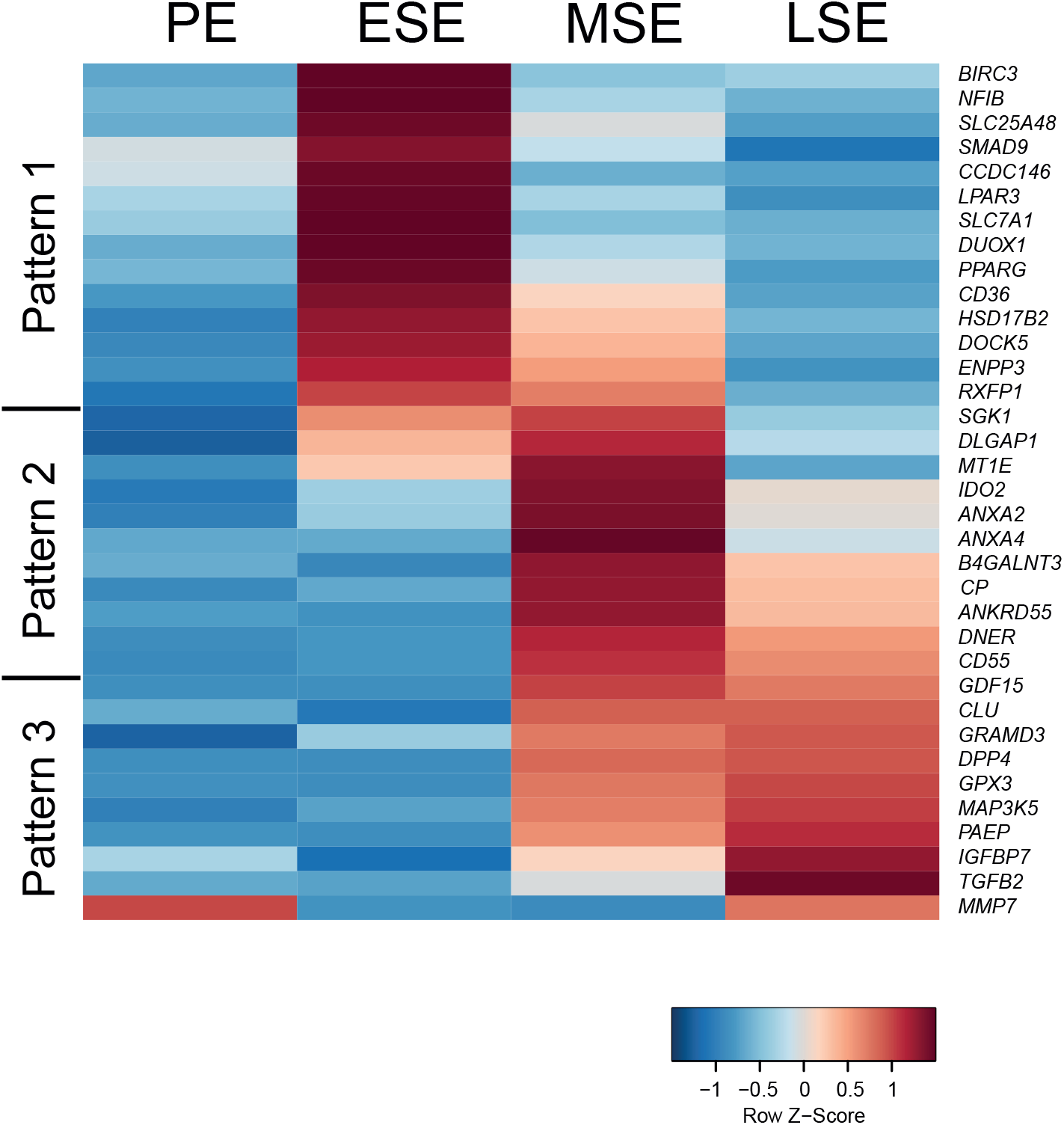
Heatmap of endometrial expression of closest gene to temporally regulated accessible chromatin loci (GEO ID: GSE4888).

**Figure S2.**
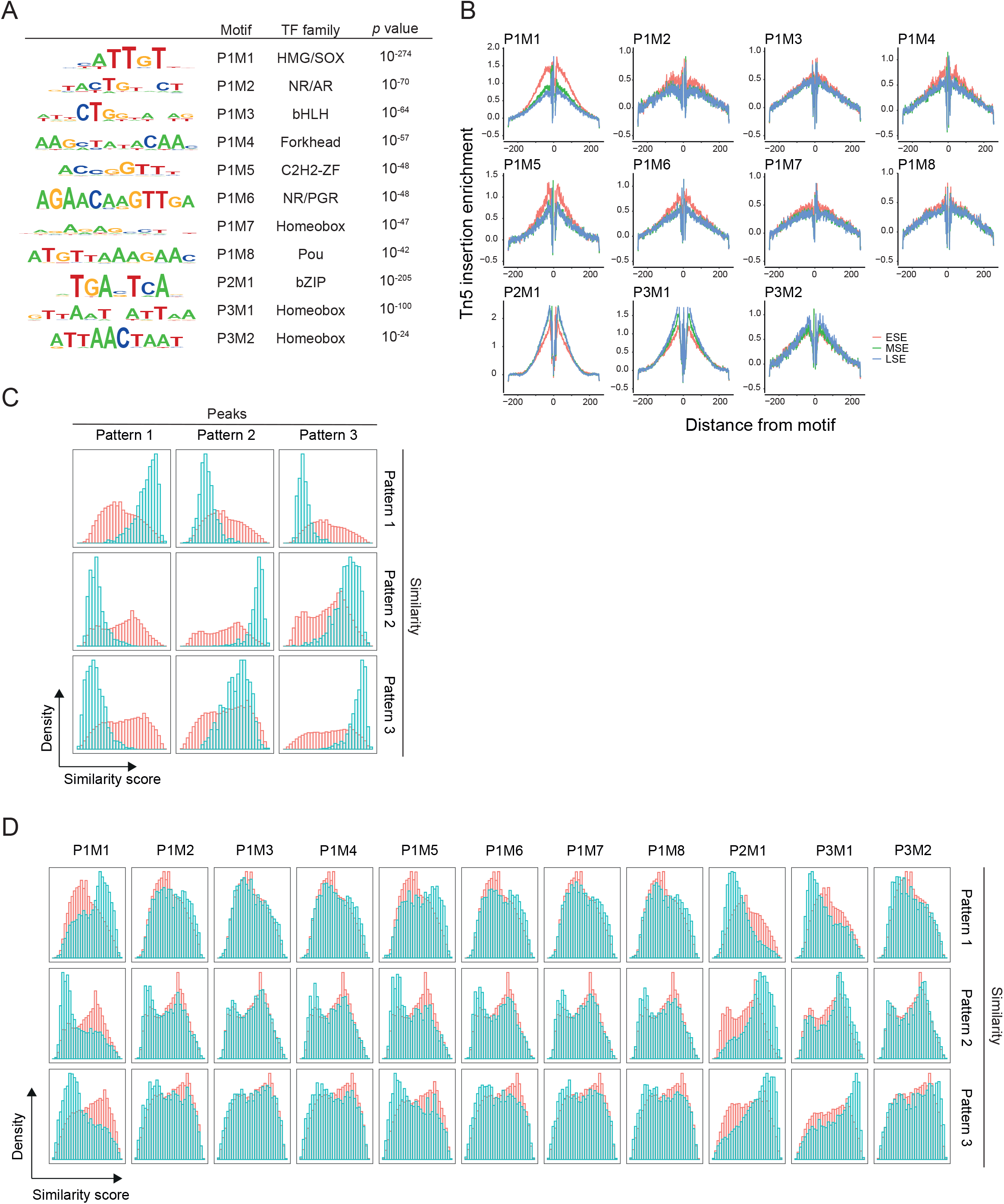
Enriched TF binding motifs in temporally regulated chromatin regions in differentiating epithelial cells. (A) Table of enriched motifs. *P* values represent binomial test results using HOMER.^65^ Motifs shown are significantly enriched based on *p* value threshold determined by performing analysis 5 times on shuffled chromatin regions. (B) Footprints analysis of temporally enriched motifs from panel A. Average transposase insertion centered around the motifs. Secretory samples were grouped as early-secretory endometrium (ESE; S18-19), mid-secretory endometrium (MSE; S20-21) or late-secretory endometrium (LSE; S22-25). (C) All ATAC-seq peaks in differentiating epithelial cells scored according to their temporal pattern similarity. Validation of pattern enrichment score in differentiating epithelial cells. (D) ATAC-seq peaks grouped by presence of enriched motifs from panel A.

**Figure S3.**
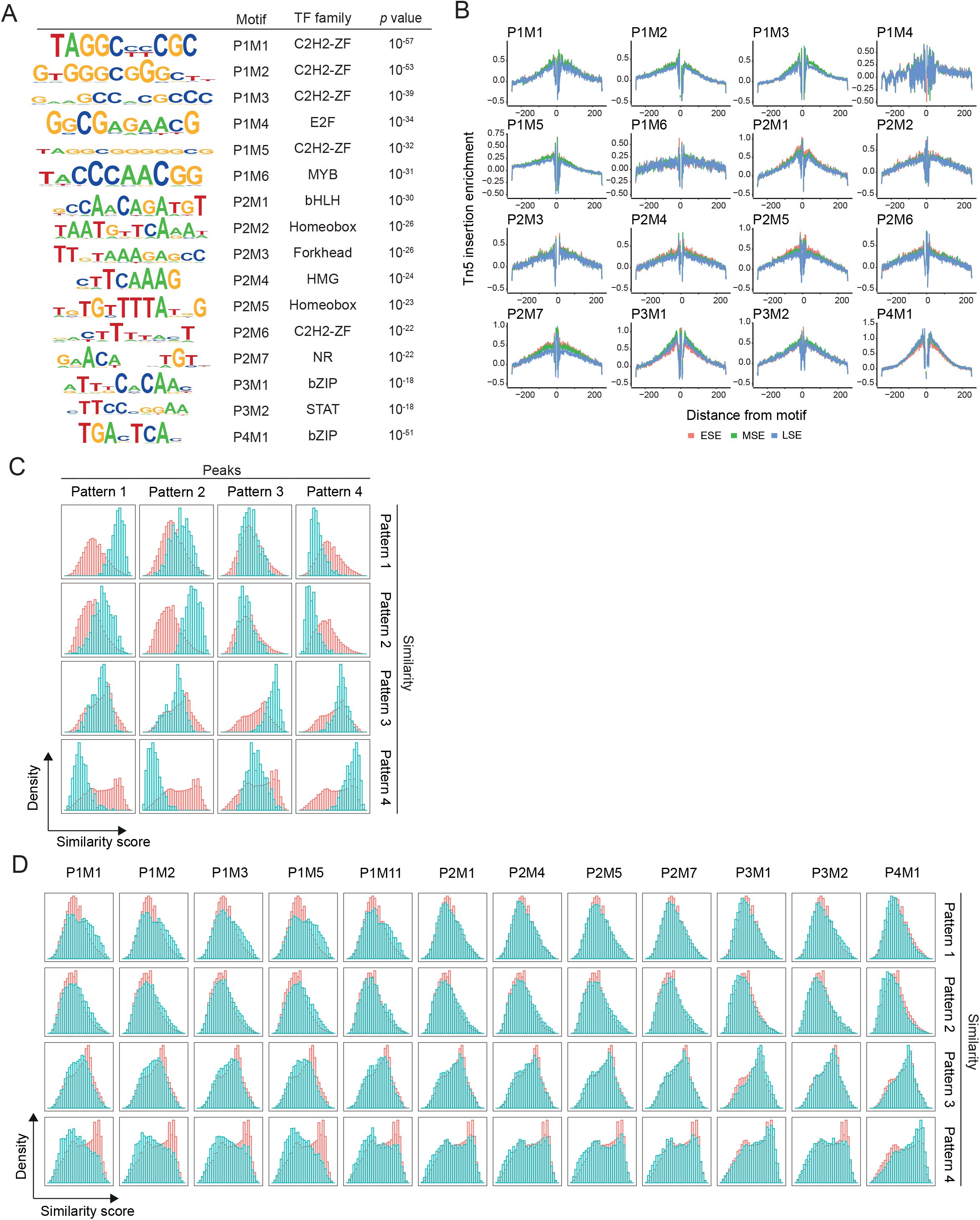
Enriched TF binding motifs in temporally regulated chromatin regions in decidualizing stromal cells. (A) Table of enriched motifs. *P* values represent binomial test results using HOMER.^65^ Motifs shown are significantly enriched based on *p* value threshold determined by performing analysis 5 times on shuffled chromatin regions. (B) Footprints analysis of temporally enriched motifs from panel A. Average transposase insertion centered around the motifs. Secretory samples were grouped as early-secretory endometrium (ESE; S18-19), mid-secretory endometrium (MSE; S20-21) or late-secretory endometrium (LSE; S22-25). (C and D) All ATAC-seq peaks in decidualizing stromal cells scored according to their temporal pattern similarity. (C) Validation of temporal pattern enrichment scores. (D) ATAC-seq peaks grouped by presence of enriched motifs from panel A.

**Figure S4.**
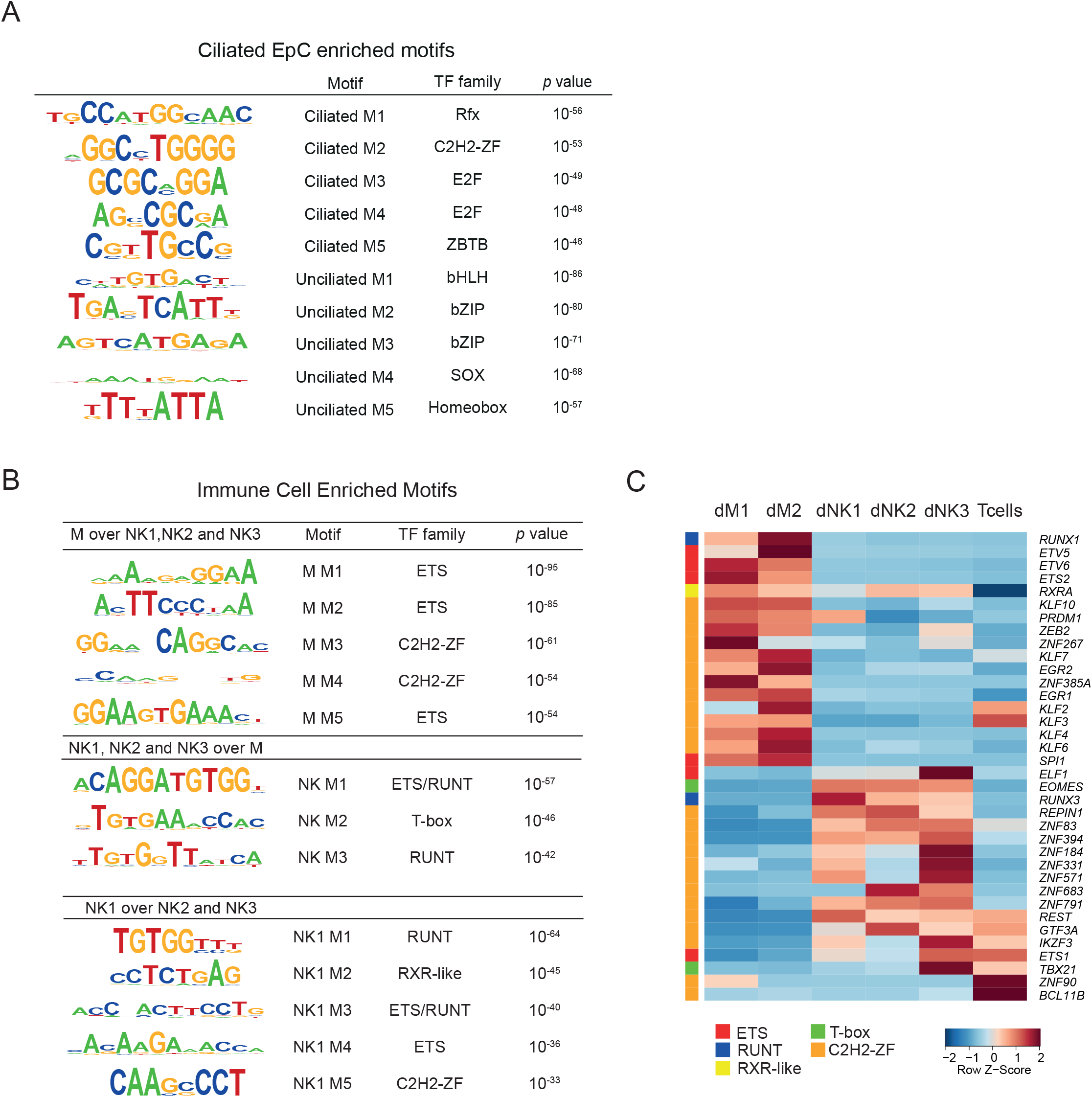
Enriched TF binding motifs in ciliated epithelial and immune cells. (A) Table of motifs enriched in ciliated and unciliated epithelial cells. (B) Table of enriched motifs in differential chromatin opening in immune cell subtypes. *P* values in represent binomial test results using HOMER.^65^ Motifs shown are significantly enriched based on *p* value threshold determined by performing analysis 5 times on shuffled chromatin regions.

**Figure S5.**
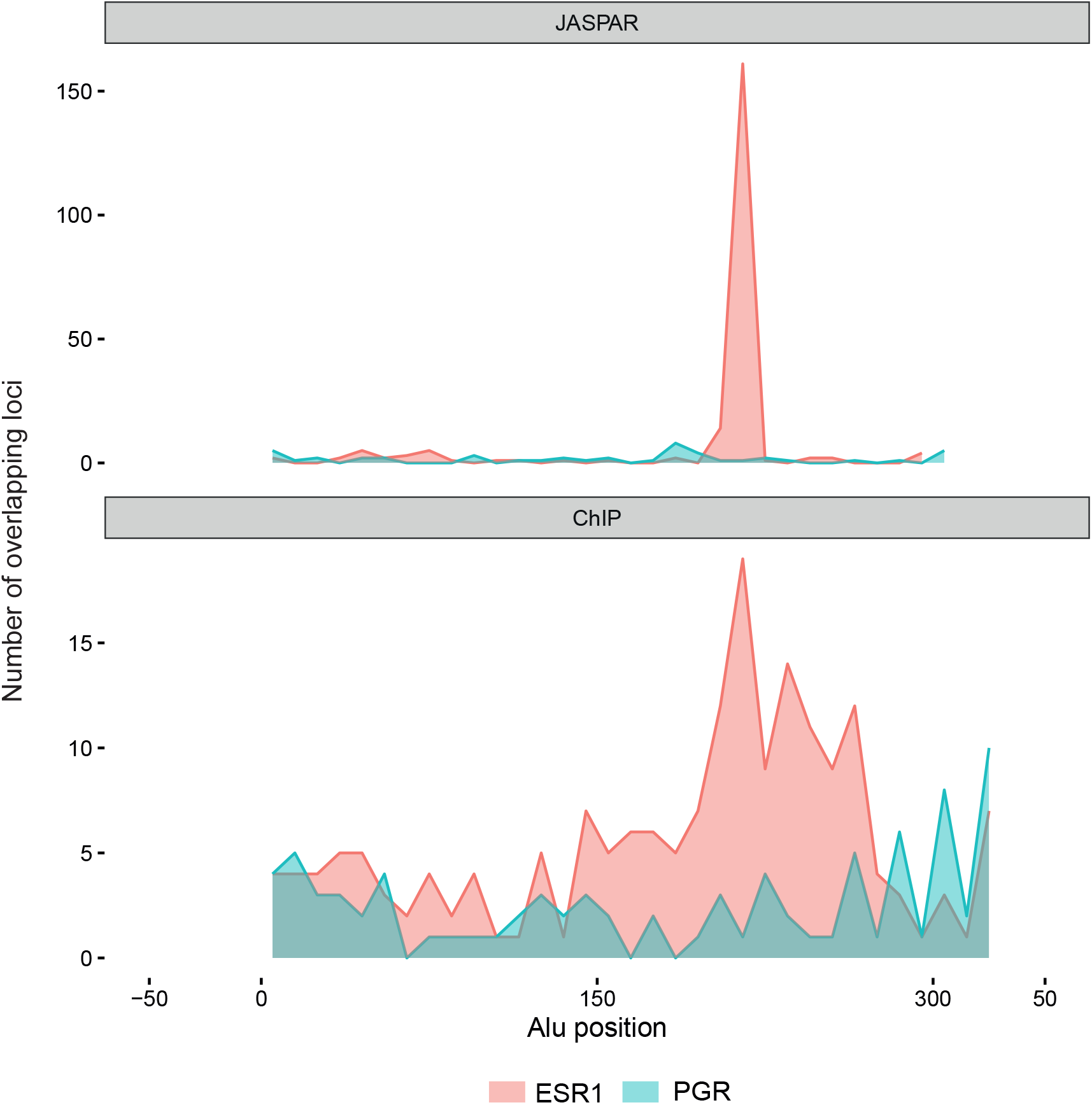
Density plots showing location of predicted and validated TFBSs from JASPAR database and ChIP-seq peak summits, respectively.

**Figure S6.**
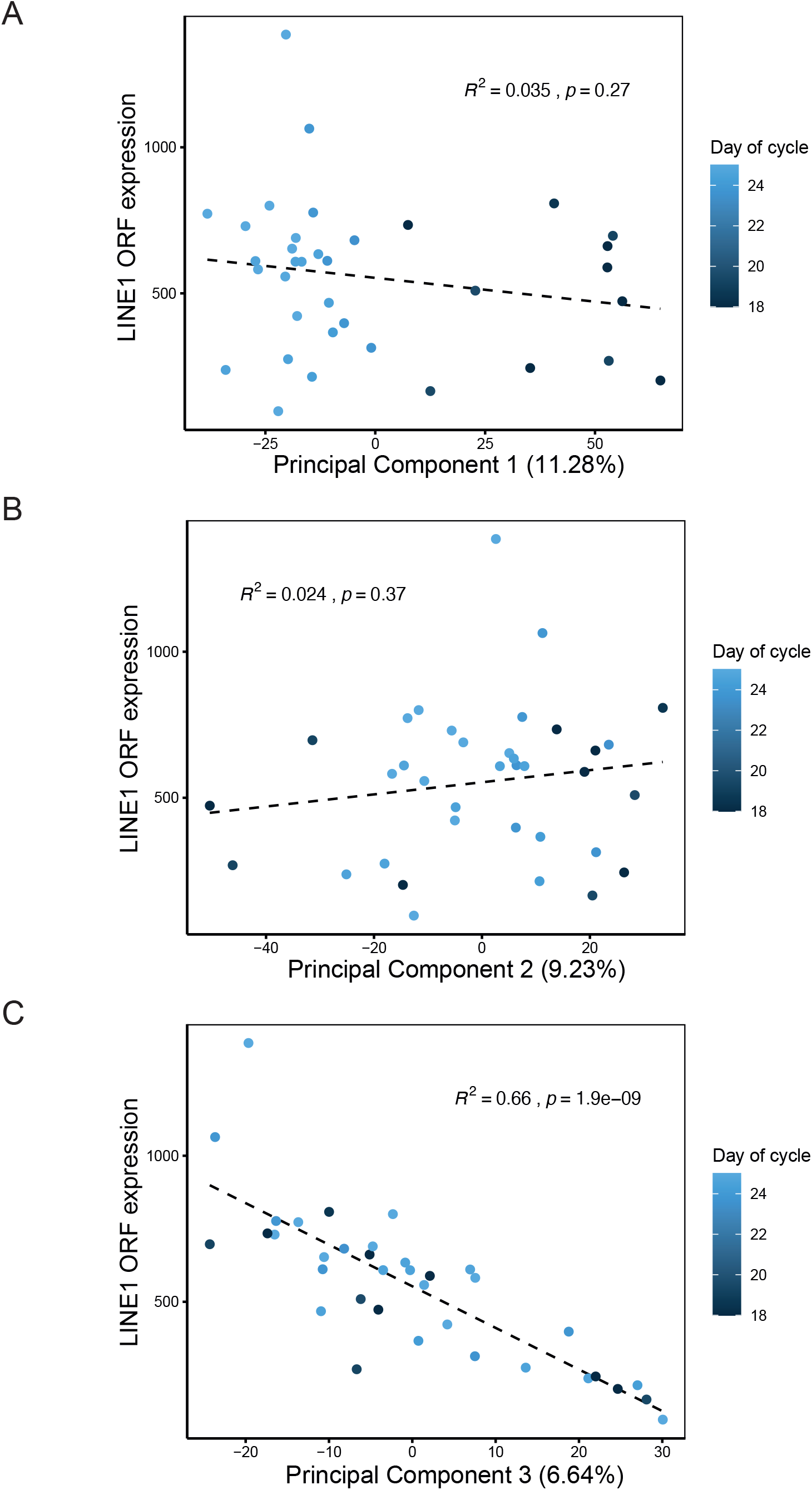
LINE1 ORF expression. Principal components in RNA-seq data obtained from 36 secretory phase endometrial samples.^18^

## STAR METHODS

### RESOURCE AVAILABILITY

#### Lead contact

Further information and requests for resources and reagents should be directed to and will be fulfilled by the lead contact, Jan Brosens.

#### Materials availability

This study did not generate new reagents.

#### Data and code availability

• The scATAC-seq data have been deposited at GEO and are publicly available as of the date of publication. Accession numbers are listed in the key resources table. This paper analyses publicly available data sets. The accession numbers for the datasets are listed in the key resources table. Immunohistochemistry data reported in this study will be shared by the lead contact upon request.

• This paper does not report original code.

• Any additional information required to reanalyze the data reported in this study is available from the lead contact upon request.

## EXPERIMENTAL MODEL AND STUDY PARTICIPANT DETAILS

The study was approved by the National Health Service (NHS) National Research Ethics­Hammersmith and Queen Charlotte’s and Chelsea Research Ethics Committee (REC reference: 1997/5065) and Tommy’s National Reproductive Health Biobank (REC reference: 18/WA/0356). Endometrial biopsies, timed relative to the last menstrual period or patient-reported pre-ovulatory luteinizing hormone surge following ovulation testing, were collected at the Implantation Research Clinic, a dedicated research clinic at University Hospitals Coventry and Warwickshire (UHCW) NHS Trust. Endometrial samples were obtained using a Wallach Endocell sampler (Wallach Surgical Devices, Trumbull, USA) and snap frozen immediately in LN2 before storage at −80 °C. Written informed consent was obtained from all participants in accordance with The Declaration of Helsinki 2000 guidelines. Anonymized endometrial biopsies were obtained from women aged between 32 and 41 years with regular cycles, body mass index between 19 and 30 kg/m2, and absence of overt uterine pathology on transvaginal ultrasound examination. Demographic details of the tissue samples are shown in Table S1.

## METHOD DETAILS

### Timing validation of secretory phase endometrial samples by RT-qPCR

Timing of secretory phase biopsies was validated by calculating the ratio of *GPX3* and *SLC15A2* gene expression as measured by RT-qPCR. RNA was extracted from endometrial biopsies which had been snap frozen in clinic (<1 min after collection), using STAT-60 (AMS Biotechnology, Oxford, UK) according to the manufacturer’s instructions. Reverse transcription was performed from 1 µg RNA using the Quantitect Reverse Transcription Kit (QIAGEN, Manchester, UK) and cDNA was diluted to 10 ng/µl equivalent before use in qPCR. Amplification was performed on a 7500 Real-Time PCR system (Applied Biosystems, Paisley, UK) in 20 µl reactions using 2 × PrecisionPlus Mastermix with SYBR Green and low ROX (PrimerDesign, Southampton, UK), with 300 nM each of forward and reverse primers. Primer sequences were as follows: *GPX3* forward: 5’-GGG GAC AAG AGA AGT CGA AGA-3’, *GPX3* reverse: 5’-GCC AGC ATA CTG CTT GAA GG-3’; *SLC15A2* forward: 5’-AGG AGG CAT CAA ACC CTG T-3’, *SLC15A2* reverse: 5’-CTA GTC CGT TCC TCT GCA TG-3’.

### Construction and sequencing of scATAC-seq libraries

Twelve timed endometrial biopsies, processed as described in detail elsewhere,^8^ were subjected to scATAC-seq. All protocols for nuclei isolation and library construction have been described previously,^67^ and available at https://support.10xgenomics.com/single-cell-atac. Briefly, nuclei were extracted according to the 10X Genomics demonstrated protocol for frozen tissue (CG000212 Rev B), modified to include an additional wash with Wash Buffer. Tissue homogenization was performed using a dounce homogenizer. Filtered nuclei suspension was pelleted as instructed and then resuspended in 50 ul Storage Buffer from the EZ nuclei kit (Sigma Aldrich, NUC101). For downstream processing, nuclei were thawed briefly on ice, resuspended with 500 ul Wash Buffer and gently pipetted. Nuclei were pelleted at 500 × g (5 min, 4 °C) then resuspended again in 500 ul Wash Buffer before counting and continuing to Chromium capture. scATAC-seq libraries were generated according to the Chromium Single Cell ATAC protocol (10X GENOMICS, CG000168) as described previously^67^ and sequenced on the Illumina NextSeq 500 platform. The sequencing data were deposited in the GEO repository (https://www.ncbi.nlm.nih.gov/geo/) under accession number GSE183771.

### Processing of scATAC-seq data

Cell Ranger ATAC pipeline version 1.2.0 was used for data pre-processing and alignment to the GRCh38 human genome assembly (refdata-cellranger-atac-GRCh38-1.2.0). When applicable, the number of fragments overlapping peaks was examined to correct the estimated number of nuclei. Peak calling was carried out on a merged BAM file using MACS2,^68^ leading to the identification of 403,434 peaks. 276,524 of these peaks were deemed high confidence (*q* < 1 x 10^−4^) and were used for further analysis. Analysis of scATAC-seq libraries, including clustering and UMAP projection was carried out with Seurat v3.2.2^69^ and Signac v0.2.5^70^ R packages. Count data was normalized using the RunTFIDF function with a scale factor of 10000. Dimensionality reduction was performed with the RunSVD function with latent semantic indexing (LSI), following which nuclei clustering and UMAP embedding was carried out with the first 50 components. The Seurat function FindAllMarkers and FindMarkers employing the Wilcoxon test was used to identify differential chromatin opening for each cell­type in the UMAP representation.

### Gene expression inference and analysis of publicly available transcriptomic datasets

Inference of gene expression levels from scATAC-seq data was performed with Signac by summing all counts from 2000 bp upstream of the transcriptional start site to the transcriptional termination site. Analysis of three endometrial scRNA-seq datasets (GEO accession GSE127918 and GSE111976, and ArrayExpress PRJEB28266)^4,7,11^ was performed using Seurat v3.2.2.^69^ The Seurat AverageExpression function was used to calculate average gene expression of published cell-types and the Seurat function FindMarkers employing the Wilcoxon test was used to identify differential gene expression. Glandular epithelial gene expression was analyzed from published RNA-seq data (GEO accession GSE84169),^66^ using the DESeq2 v1.30.1 package in R. Expression profiling by array of human endometrium (GEO accession GSE4888)^22^ was visualized with heatmap function of gplots v3.1.3 package in R. Gene Ontology enrichment analysis was performed with PANTHER.^71^

### Temporal analysis of scATAC-seq peaks and similarity scores

To perform temporal pattern analysis of chromatin changes in secretory phase samples, scATAC-seq peaks with temporal variation were first selected by pairwise comparisons. Peaks with at least one significant change between secretory samples were then grouped into temporal patterns by K-means clustering on scaled averaged values. To measure the likelihood of non-clustered peaks fitting the secretory phase temporal patterns, we devised similarity scores. At every timepoint, the average opening of the peak was compared against the mean and standard deviation obtained from all the peaks in a temporal pattern to produce a z-score. Finally, an overall similarity score was obtained by adding the absolute value of the individual z-scores and subtracting this value from the maximum such that higher similarity scores represented peaks with a pattern close to the original (Figures S2C and S3C).

### TF binding motif analyses

Differentially opening chromatin regions were interrogated for enriched short sequence motifs using HOMER v4.8.^65^ To establish the significance cut-off for enriched motifs, the analysis was repeated 5 times on shuffled data. Footprint and PlotFootprint functions in Signac were used to visualise the average chromatin opening pattern around the identified motifs. Chromatin opening at a specific site was visualised with the CoveragePlot function in Signac and adapted using the ggplot2 R package. To help predict the specific transcription factors responsible for the observed chromatin changes, every TF from the enriched families were first identified using AnimalTFDB v3,^72^ then examined RNA expression of members of TF families identified.^4^ For ESR and PGR binding site enrichment analysis, the entire genome was scanned for ERE and PRE motifs using findmotifs function of HOMER. Peaks were deemed to contain ERE or PRE sites if they overlapped motifs (1bp).

### Transposable elements

Genomic coordinates for repetitive elements were obtained from the RepeatMasker track of the UCSC genome browser (A.F.A. Smit, R. Hubley & P. Green RepeatMasker at http://repeatmasker.org). Enrichment of repeats overlapping temporal chromatin opening patterns was calculated using the hypergeometric distribution in R. Full length Alu sequences were analyzed for evidence of transposase accessibility. TFBSs predictions for ESR1, FOSL2, GATA2, FOXO1, and CEBPB were obtained from JASPAR track in UCSC browser.^73^ TF binding data were obtained from published ChIP-seq datasets.^31–35^ Peak summits were generated with MACS2^68^ on sequencing files mapped to the hg38 genome assembly using STAR aligner.^74^ For spatial mapping of TE-derived transcripts in the endometrium and adjacent myometrium, we reanalyzed publicly available Visium (10X GENOMICS) data (ArrayExpress ID: E-MTAB-9260; 152807).^37^ Locus-specific quantification of TE-derived transcripts in the spatial transcriptomic data was performed with SoloTE.^38^ The resulting count matrix was used for single-cell analysis using Seurat. Visualization was performed by aggregating the normalized expression of TE families. Locus-specific quantification of L1 ORF expression in bulk RNA-seq data (GEO accession GSE180485) was performed with L1EM.^75^

### HNF1B immunohistochemistry

Endometrial biopsies were fixed overnight in 10% neutral buffered formalin at 4°C and wax embedded in Surgipath Formula ‘R’ paraffin (Leica BioSystems, Wetzlar, Germany) using the Shandon Excelsior ES Tissue processor (ThermoFisher Scientific, Massachusetts, US). Tissues were sliced into 3 μm sections using a microtome and adhered to coverslips by overnight incubation at 60°C. Deparaffinization, antigen retrieval (sodium citrate buffer; 10 mM sodium citrate, 0.05% Tween-20, pH 6), antibody staining, haematoxylin counter stain and DAB colour development were fully automated in a Leica BondMax autostainer (Leica BioSystems, Wetzlar, Germany). Tissue sections were labelled for HNF1B (HPA002083, Prestige Antibodies_®_ Powered by Atlas Antibodies, Sigma-Aldrich, Missouri, US) using a 1:1000 dilution. Stained slides were de-hydrated, cleared and cover-slipped in a Tissue-Tek Prisma Automated Slide Stainer, model 6134 (Sakura Flinetek, Inc., California, US) using DPX coverslip mountant. Bright-field images were obtained on a Mirax Midi slide scanner with a 20× objective lens and viewed in CaseViewer v2.4.0.1198028 (3DHISTECH Ltd, Budapest, Hungary). Three randomly selected areas of interest underlying the luminal epithelium were captured and each of the regions was divided manually into three compartments: stroma, glandular epithelium, and luminal epithelium. ImageJ image analysis software (Rasband, W.S., ImageJ, National Institutes of Health) was used to quantify HNF1B^+^ cells in the glandular epithelium with staining intensity manually determined by background thresholding. The percentage of HNF1B^+^ cells was calculated (i.e., HNF1B^+^ cells/total cells ×100) in each region of view for the glandular epithelium.

## Notes

### Competing Interest Statement

The authors have declared no competing interest.

